# Mitochondrial Bioenergetic Failure in SLE Immunocytes: Targeting Fitness for Therapy

**DOI:** 10.1101/2025.06.10.658845

**Authors:** Anjali S. Yennemadi, Natasha Jordan, Sophie Diong, Mark Little, Joseph Keane, Gina Leisching

**Affiliations:** Department of Clinical Medicine, Trinity Translational Medicine Institute, St James’s Hospital, Trinity College Dublin, The University of Dublin, Dublin, Ireland; Department of Rheumatology, St. James’s Hospital, James Street, Dublin, Ireland; Department of Dermatology, St. James’s Hospital, James Street, Dublin, Ireland; Department of Nephrology, St. James’s Hospital, James Street, Dublin, Ireland; Trinity Kidney Centre

**Keywords:** Systemic Lupus Erythematosus, Mitochondrial dysfunction, CD4+, T cells, CD8+, T cells, B cells, Oxidative phosphorylation, Metabolic therapy

## Abstract

**Background:** Systemic Lupus Erythematosus (SLE) is characterized by dysregulated immune responses linked to immunometabolic perturbations. While mitochondrial dysfunction has been implicated in SLE, its cell-type-specific impact on immune subsets remains underexplored.

**Methods:** We repurposed existing RNA-seq data from SLE patient peripheral blood mononuclear cells, with a focus on nuclear-encoded mitochondrial (NEmt) genes, as well as mitochondrial genes themselves, to identify differentially expressed genes compared to healthy controls. Mitochondrial stress tests were performed on freshly isolated CD4+ T cells, CD8+ T cells, B cells, and monocytes from SLE patients and healthy donors to assess bioenergetic function.

**Results:** RNA-seq revealed that both NEmt genes and mitochondrial genes were downregulated in the PBMC population of SLE patients. In situ mitochondrial stress tests revealed significant reductions in oxygen consumption rate (OCR), indicating impaired oxidative phosphorylation (OXPHOS) across all immune subsets, while extracellular acidification rate (ECAR), a marker of glycolysis, remained unchanged. These findings highlight immune-cell-specific mitochondrial bioenergetic failure in SLE, without compensatory glycolytic adaptation.

**Conclusion:** Our results position mitochondrial fitness as a novel therapeutic target in SLE. We propose leveraging high-throughput screening of mitochondria-targeted compounds, including FDA-approved agents, to enhance OXPHOS, regulate mitophagy, or mitigate oxidative stress.

This precision-based approach offers a paradigm shift from conventional immunosuppression to metabolic recalibration, with the potential to restore immune homeostasis in SLE.

**Key messages:**

- **What is already known on this topic:** Mitochondrial dysfunction and metabolic reprogramming have been linked to SLE pathogenesis, but the cell-type-specific extent of mitochondrial impairment and its therapeutic potential remained unclear.
- **What this study adds:** We demonstrate reduced OXPHOS in CD4+ T cells, CD8+ T cells, B cells, and monocytes from SLE patients for the first time, highlighting cell-type-specific mitochondrial dysfunction as a possible driver of immune dysregulation and target for therapy.
- **How this study might affect research, practice, or policy:** This study advocates for drug repurposing to restore mitochondrial fitness, offering a novel therapeutic strategy to complement existing SLE treatments and improve patient outcomes.

## Introduction

Systemic lupus erythematosus (SLE) is a chronic autoimmune disorder characterized by dysregulated immune responses, including hyperactivity of B cells and loss of tolerance in T cells. While genetic and environmental factors contribute to SLE pathogenesis, emerging evidence highlights metabolic dysregulation as a critical driver of immune dysfunction^1^.

Mitochondria, beyond their role as a crucial source of ATP, are central to immune cell homeostasis, influencing activation, differentiation, and apoptosis through oxidative phosphorylation (OXPHOS), reactive oxygen species (ROS) signaling, and metabolic intermediates. More than 99% of mitochondrial proteins are encoded by nuclear DNA and therefore also play a significant role in mitochondrial function in addition to mitochondrial genes themselves ^2^. Mitochondria have been described as a key player in SLE and targeting these organelles may be a worthwhile therapeutic option^3^. Notably, CD4+ and CD8+ T cells rely on mitochondrial fitness for effector function and memory formation, B cells require mitochondrial activity for plasma cell survival, and mitochondria in monocytes play a central role in regulating polarisation and differentiation^4^. Despite growing interest in metabolic reprogramming in autoimmunity research, the role of mitochondrial dysfunction in specific immune subsets in SLE remains poorly defined.

This study aimed to highlight mitochondrial metabolic dysfunction in immune cell subsets of SLE patients by combining repurposed RNA-seq data with real-time *in situ* bioenergetic analysis. By integrating these approaches, we sought to emphasize the critical role of mitochondrial fitness in immune cell homeostasis and introduce the narrative that restoring mitochondrial function through existing drugs represents a promising therapeutic strategy. Targeting mitochondrial bioenergetics could restore immune cell functionality, offering a novel approach to ameliorating disease severity in SLE.

## Materials and Methods

Using an existing bulk RNA-seq dataset of PBMCs ^5^ we randomly selected healthy controls (HC) and SLE patients (Table S1) to characterise the differential expression of genes associated with mitochondrial metabolism and function. Read quality was assessed and reads were mapped to the human reference genome (GRCh38/hg38) using STAR (v2.7.4a). Raw counts were generated using HTSeq (v2.0.1), and differential expression (DE) analysis was carried out using DESeq2 (v1.36.0). Gene lists were compiled using KEGG, and MitoCarta 2.0 release 2015 ^6^and imported into in Rstudio (R v4.3.3) using Bioconductor (v3.15). Data visualisation was performed using packages EnhancedVolcano (v1.10.0), ggplot2 (v3.4.0), pheatmap (v1.0.12), and ggrepel (v0.9.2).

Ethical clearance for blood collection was obtained from St James’s Hospital/Tallaght University Hospital Joint Research Ethics Committee. For functional metabolic work, blood was collected from consented SLE patients at St. James’s Hospital, Dublin as well as sex-matched healthy controls (Table 1). PBMCs were isolated using Lymphoprep (STEMCELL Technologies) whereafter CD4+ T cells, CD8+ T cells, B cells and monocytes were isolated using a magnetic bead-based negative selection method (STEMCELL Technologies). Cells were counted and plated into a Seahorse XF24 cell culture microplate (Agilent Technologies, Inc.) and mitochondrial function was assessed using the Cell MitoStress Test (Agilent Technologies, Inc.) on the day of isolation. All values were normalized using the Crystal Violet dye extraction growth assay and the Agilent Wave Desktop 2.6 Software (https://www.agilent.com) was used for analysis.

**Table 1.** Demographics and clinical characteristics of HC and patients with lupus examined in this study.

| Characteristics | Lupus (N= 7) | HC (N= 7) |
| --- | --- | --- |
| **<u>Demographics</u>** |  |  |
| Age median, years (range) | 53 (23 - 62) | 36 (24 - 68) |
| Sex N (%) |  |  |
| Female | 6 (85.71%) | 6 (85.71%) |
| Male | 1 (14.29%) | 1 (14.29%) |
| **<u>Lupus History</u>** |  |  |
| Disease Severity |  | - |
| SLEDAI > 4 | 3 (42.85%) | - |
| SLEDAI ≤ 4 | 4 (57.14%) | - |
| **<u>Medications</u>** |  |  |
| Hydroxychloroquine | 0 (0%) | - |
| MMF | 4 (57.14%) | - |
| Mepacrine | 3 (33.33%) | - |
| Azathioprine | 0 (0%) | - |
| Methotrexate | 2 (28.57%) | - |
| Plaquenil | 4 (57.14%) | - |
| Prednisolone | 4 (57.14%) | - |
| Tacrolimus | 1 (11.11%) | - |
| Warfarin | 0 (0%) | - |
| Valsartan | 1 (11.11%) | - |
| Aspirin | 1 (11.11%) | - |
| Topical betnovate | 1 (11.11%) | - |

## Results

RNA-seq analysis revealed 61,544 DE genes between SLE and HC PBMCs (Fig. 1A), of which 9953 were significantly DE genes (Tables S2-S5). As the focus of our study was to explore the extent of mitochondrial metabolic dysfunction between SLE and HC PBMCs, we limited gene annotation and defined the following lists for further analysis: (i) nuclear-encoded mitochondrial (NEmt) genes – that are encoded in the nucleus, and code for proteins that are imported into as well as function within the mitochondria (Fig.1B); (ii) mitochondrial (mt) genes – genes that are encoded within mitochondrial DNA (Fig. 1C); and (iii) mt-associated genes – a combined list of NEmt and mt genes (Fig. 1D).

**Figure 1.**
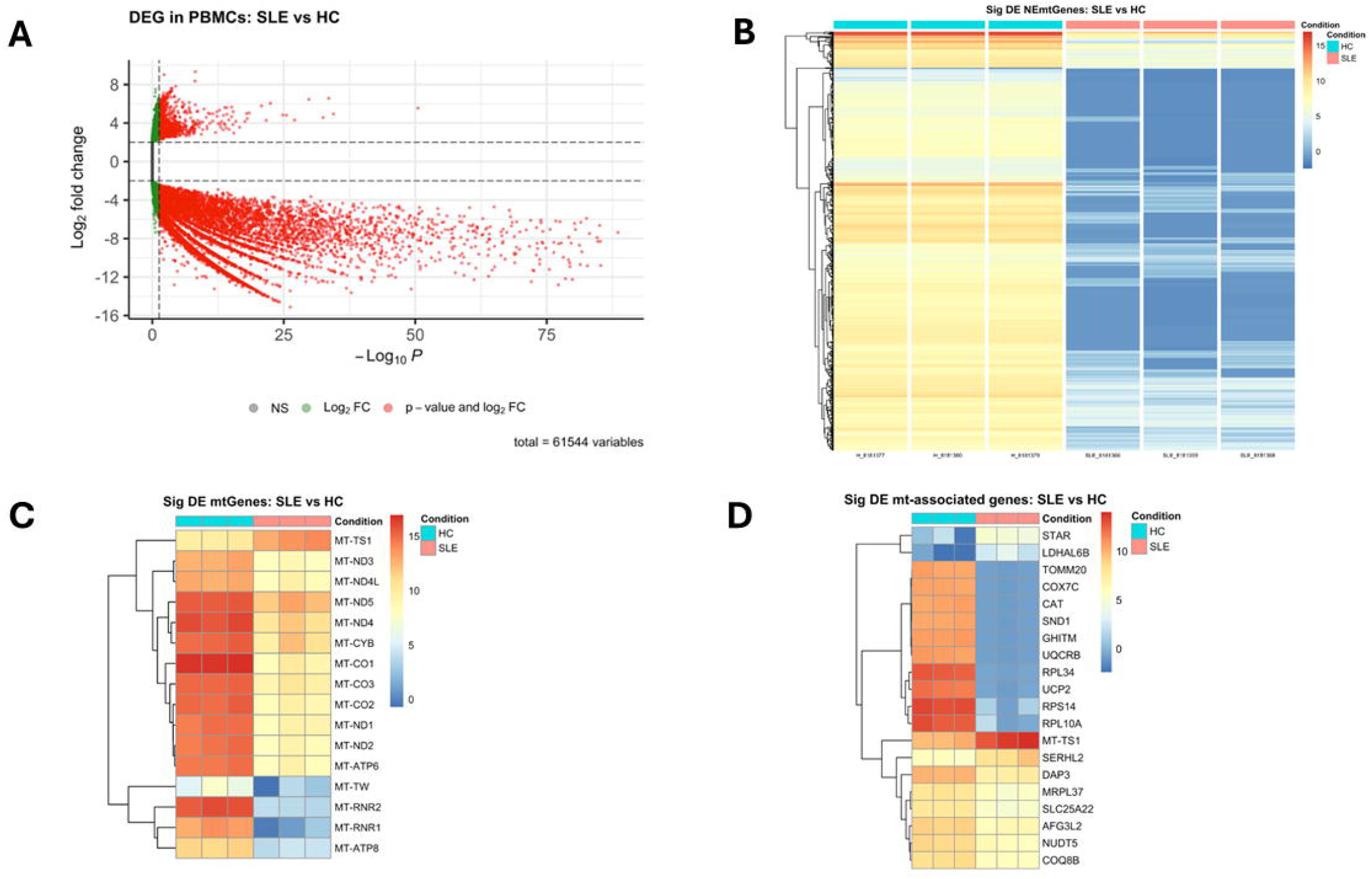
RNA-seq analysis of existing data reveal global downregulation of nuclear-encoded mitochondrial genes and mitochondrial-encoded genes in PBMCs from SLE patients. A. RNA-seq analysis revealed 61,544 DE genes between SLE and HC PBMCs. B. Heatmap of DE NEmt genes shows global downregulation of genes in SLE PBMCs. C, D. Heatmap representing the top up-and downregulated mt genes and mt-associated in SLE patients vs. HC. Yellow-red=upregulated, blue=downregulated, padj <0.05, log2FC >|2|).

We observe a significant overall downregulation of NEmt genes in SLE patients (padj <0.05, log2FC >|2|Fig. 1B). Only 15 differentially expressed mt genes displayed significant downregulation, except for *MT-TS1*, compared to HC. Most notably, the MT-ND family of genes responsible for encoding NADH dehydrogenase (complex I), MT-ATP6 encoding mitochondrial complex V (Fig.1C), and MT-CYB and MT-CO1-3 encoding complex II, exhibit downregulation compared to healthy controls. Fig. 1D represents the top 20 most up-and down-regulated mt-associated genes. Of note, genes essential for protein-transporting ATPase activity (*TOMM20*), mitochondrial electron transport chain (*COX7C*), mitophagy (*SND1*), mitochondrial calcium:proton antiporter activity (GHITM), and the regulator of mitochondrial proton leak (UCP2) were all downregulated (Fig. 1D).

In line with the above data, the Cell MitoStress test revealed striking functional mitochondrial defects in all immune subsets from SLE patients compared to their sex-matched HC (Fig.2A-D). Lower basal OCRs and unchanged ECARs were observed across all cell types, as well as abnormal oxygen consumption curves, suggesting deficiencies in mitochondrial metabolism and function only, with no glycolytic compensation.

**Figure 2.**
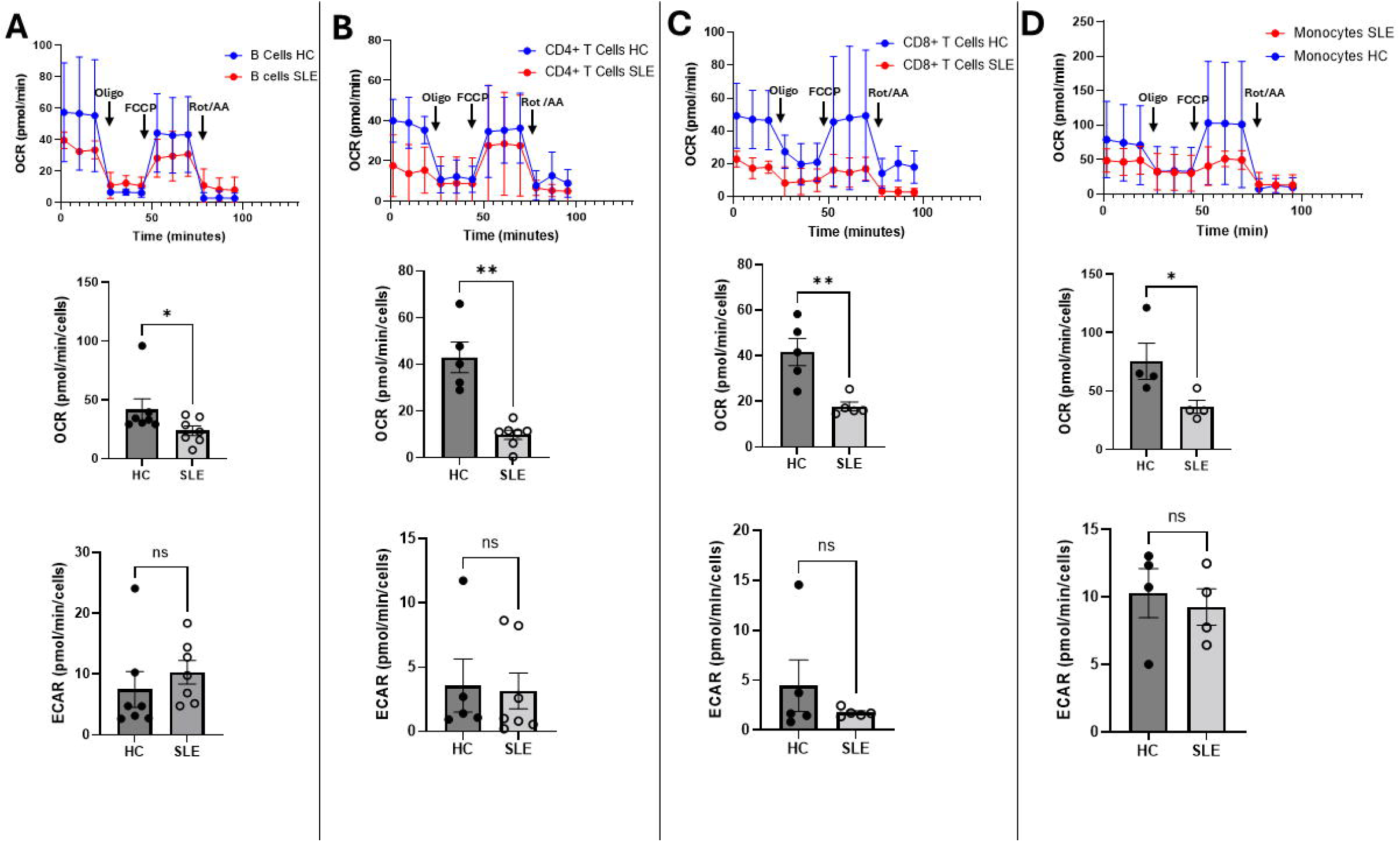
B cells, CD4+ T cells, CD8+ T cells, and monocytes from SLE patients exhibit reduced OCR with unchanged ECAR compared to healthy controls. The Agilent Mitostress test was used to assess mitochondrial fitness and function during sequential treatment with oligomycin, carbonyl cyanide-4-(trifluoromethoxy)phenylhydrazone (FCCP) and rotenone + antimycin A (Rot/AA). Decreased basal OCR and atypical Mitostress profiles in **A**. B cells, **B**. CD4+ T cells, **C**. CD8+ T cells and **D**. monocytes from patients with SLE. ECAR remains unchanged across all subtypes. Data represented as mean +-SEM. Mann-Whitney U-test, n=4-7 HC and n=4-7 SLE.

## Discussion

Our study provides compelling evidence of widespread mitochondrial dysfunction across multiple immune cell populations in SLE patients. Through direct bioenergetic profiling, we demonstrate impaired oxidative phosphorylation in CD4+ T cells, CD8+ T cells, B cells, and monocytes, and establish a pattern of pan-immune metabolic disruption that distinguishes human SLE from murine models where OXPHOS upregulation is typically observed^7 8^. The unchanged glycolytic activity despite OXPHOS deficiency suggests these cells lack the metabolic flexibility to compensate for mitochondrial impairment, potentially explaining the functional deficits observed in SLE immune cells. Our RNA-seq analysis further pinpoints complexes I, II and V as particularly vulnerable components of the respiratory chain in PBMCs from SLE patients.

These findings open two promising therapeutic avenues that warrant further investigation. First, comprehensive respiratory chain complex testing could identify specific electron transport chain vulnerabilities to guide targeted interventions^9^. Our transcriptomic findings suggest the downregulation of several mitochondrial complexes, however it is unclear in which immune cells these deficiencies are occurring. While several scRNAseq studies have helped to unravel the transcriptomic landscape of SLE PBMCs, there is still very limited work exploring the immunometabolism of SLE PBMCs. Indeed, scRNAseq studies avoid performing mitochondrial gene expression assessments as a high abundance of mitochondrial transcripts is often used as an indicator of poor sequencing quality or cell death, and such cells are excluded from analysis^10^. Thus, although our study provides a relatively limited overview, it remains one of very few to exclusively explore differential expression across mitochondrial and NEmt genes in SLE PBMCs. Concurrent research need for immunocyte -specific mitochondrial gene assessment is also required to aid this therapeutic approach with more focussed target generation allowing for better precision.

Second, the growing availability of mitochondria-targeted compound libraries, including FDA-approved agents, presents immediate opportunities for drug repurposing to address OXPHOS deficiencies, regulate mitophagy, or reduce oxidative stress. A study using this method discovered that etravirine could be a promising treatment for Friedreich’s ataxia. The research found that etravirine restores the function of the iron-sulfur cluster enzyme aconitase and improves oxidative stress resistance in cells taken from Friedreich’s ataxia patients ^11^. Drug repurposing therefore holds particular promise given the established safety profiles of many candidate compounds. Additionally, the use of mitochondriotropics as well as therapeutic strategies applied to patients with inborn errors of metabolism or inherited mitochondrial diseases may also present a potential avenue to restore mitochondrial function^12^.

In conclusion, our findings highlight cell-type-specific mitochondrial dysfunction as a central feature of SLE dysregulated immune cell metabolism, with impaired OXPHOS in CD4+ T cells, CD8+ T cells, B cells and monocytes likely contributing to immune dysregulation. While previous studies have broadly proposed targeting mitochondria in SLE, our data underscore the need for precision in therapeutic strategies. By focusing on agents that selectively address the bioenergetic deficits observed in our study, we can move beyond generalized immunosuppression toward tailored metabolic therapies. Future studies should prioritize functional validation of these compounds in preclinical SLE models, with the ultimate goal of translating mitochondrial restoration into clinical benefit for patients. Prioritizing mitochondrial health may potentiate novel therapies to recalibrate immune homeostasis in SLE.

## Acknowledgments

We would like to thank Dr. Lorraine Thong and Dr. Kevin Brown for their assistance in collecting samples from healthy donors for this manuscript.

## References

1. Yennemadi AS, Keane J, Leisching G. Mitochondrial bioenergetic changes in systemic lupus erythematosus immune cell subsets: Contributions to pathogenesis and clinical applications. Lupus 2023;32(5):603–11.

2. Ferramosca A, Zara V. Biogenesis of mitochondrial carrier proteins: molecular mechanisms of import into mitochondria. Biochimica Et Biophysica Acta (BBA)-Molecular Cell Research 2013;1833(3):494–502.

3. Quintero-González DC, Muñoz-Urbano M, Vásquez G. Mitochondria as a key player in systemic lupus erythematosus. Autoimmunity 2022;55(8):497–505.

4. Ravi S, Mitchell T, Kramer PA, et al. Mitochondria in monocytes and macrophages-implications for translational and basic research. The International Journal of Biochemistry & Cell Biology 2014;53:202–07. doi: 10.1016/j.biocel.2014.05.019

5. Tokuyama M, Kong Y, Song E, et al. ERVmap analysis reveals genome-wide transcription of human endogenous retroviruses. Proceedings of the National Academy of Sciences 2018;115(50):12565–72.

6. Calvo SE, Clauser KR, Mootha VK. MitoCarta2. 0: an updated inventory of mammalian mitochondrial proteins. Nucleic acids research 2016;44(D1):D1251–D57.

7. Yin Y, Choi S-C, Xu Z, et al. Normalization of CD4+ T cell metabolism reverses lupus. Science translational medicine 2015;7(274):274ra18–74ra18.

8. Yin Y, Choi S-C, Xu Z, et al. Glucose oxidation is critical for CD4+ T cell activation in a mouse model of systemic lupus erythematosus. The Journal of Immunology 2016;196(1):80–90.

9. Salabei JK, Gibb AA, Hill BG. Comprehensive measurement of respiratory activity in permeabilized cells using extracellular flux analysis. Nature protocols 2014;9(2):421–38.

10. Yates J, Kraft A, Boeva V. Filtering cells with high mitochondrial content depletes viable metabolically altered malignant cell populations in cancer single-cell studies. Genome Biology 2025;26(1):91.

11. Alfedi G, Luffarelli R, Condò I, et al. Drug repositioning screening identifies etravirine as a potential therapeutic for friedreich’s ataxia. Movement Disorders 2019;34(3):323–34.

12. Andreux PA, Houtkooper RH, Auwerx J. Pharmacological approaches to restore mitochondrial function. Nature reviews Drug discovery 2013;12(6):465–83.

